# The impact of epistasis on the genomic prediction of non-assessed and non-overlapping single crosses in multi-environment trials

**DOI:** 10.1101/2025.02.21.639587

**Authors:** José Marcelo Soriano Viana, Jean Paulo Aparecido da Silva

## Abstract

Because no previous simulation-based study assessed the influence of epistasis on the genomic prediction of untested single crosses (SCs), the objective was to assess the impact of epistasis on the genomic prediction of untested SCs in multi-environment trials (METs), assuming seven types of digenic epistasis. We genotyped two groups of 70 doubled-haploid lines and phenotyped all 4900 SCs in five environments. The average density for SNPs was 0.03 cM. Regarding the distribution of the SCs over environments, we adopted 10 and 30% of tested SCs and 80% of non-overlapping SCs. To assess the efficacy of the genomic prediction we computed coincidence index (CI) and prediction accuracy. The percentage of the genotypic variance due to epistasis ranged from 18 to 48%. The CI ranged from 0.24 to 0.38, under the lower training set, and between 0.30 to 0.50, assuming the higher training set. Fixing complementary epistasis and increasing the ratio epistatic variance/genotypic variance from 18 to 39% led to a decrease in the coincidence index in the range from 6.5 to 22.0%. Accuracy showed a positive correlation with CI. Assuming epistasis but fitting the additive-dominance model led to a decrease in the CI. Epistasis can significantly affect the genomic prediction of SCs in METs, depending on type, proportion of the genotypic variance due to epistasis. The prediction accuracy efficiently expresses the efficacy of genomic prediction of untested SCs. The genomic prediction of untested single crosses in each environment and across environments is very effective, even under a low training set size.

## Introduction

The most significant application of genomic selection in plant breeding is the prediction of non-assessed hybrids or doubled haploid (DH)/inbred/pure lines in multi-environment trials (METs). The significance is even higher for maize hybrid breeding. Since the development of the doubled haploid technology, maize seed companies generate a huge number of homozygous genotypes in a single year from proprietary superior single crosses or biparental populations, by a fraction of the cost to develop the same number of inbred lines from several generations of selfing (Chaikam *et al*. 2019). Thus, the costs and time required to develop maize homozygous genotypes decreased but the challenge of identifying the best single crosses remains because they need to be extensively assessed. Genotyping a limited number of maize DH/inbred lines for thousands of single nucleotide polymorphisms (SNPs) and using a sparse testing system to distribute a sample of the possible single crosses in a limited number of plots in METs can allow, by genomic prediction of complex traits, at least an initial evaluation of thousands of non-assessed single crosses (Jarquin *et al*. 2020). In the next stage, the breeder can use his limited resources to field assessment of a reduced number of superior predicted single crosses. Furthermore, because the genomic prediction model includes a genotype x environment (GxE) interaction effect, breeders can optimize the allocation of their superior predicted single crosses in specific environments.

Breeders agree that fitting GxE effect in METs increases the prediction accuracy. Using a wheat line panel and climatic covariates for the environment characterization, Robert *et al*. (2020) assessed the prediction accuracy of untested genotypes and of new environment for grain yield and observed average increases of 50 and 30%, respectively, by including the GxE effect. Martini *et al*. (2020) assessed two covariance structures for GxE interactions using wheat- and maize-panels, both assessed in four environments. They theoretically demonstrated that there is no difference between the Hadamard and Kronecker formulations for the covariance matrix of GxE effects. The authors assumed 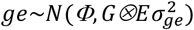 and *e*∼*N*(Φ, *Eσ*^2^), where *G* is the additive genomic relationship matrix and *E* is the error correlation matrix. They also assumed *E* = *I* and *E* estimated by the pure/inbred lines phenotypic correlations between environments. Compared to the yield accuracies within each environment (no GxE effect), the inclusion of the GxE effects increased the prediction accuracy in each environment in 75% of the scenarios. The increases ranged from 1 to 38% (11% on average). Processing zinc concentration of three maize panels assessed in three environments, Mageto *et al*. (2020) observed higher accuracies for sparse testing than for prediction of newly developed lines, genetically related with those in the training set. The prediction accuracy increases by including GxE effects, under sparse testing, were not significant. The use of sparse testing reduces the costs of assessing a high number of genotypes by optimizing the allocation design using genome-based models with GxE interaction (Jarquin *et al*. 2020). Higher prediction accuracy of non-assessed maize single crosses in METs can be achieved by combining non-linear kernel methods, dominance, and envirotyping data (up to 85% compared to genomic-BLUP) (Costa-Neto *et al*. 2020).

Regarding the significance of non-additive effects for genomic prediction, modelling dominance effects for the prediction of grain yield of non-assessed maize single crosses can significantly increase the accuracy in METs (Ferrao *et al*. 2020). Because epistasis can significantly affect heterosis, maize breeders expect the same positive impact on the grain yield prediction accuracy by additionally modelling epistatic effects. Roth *et al*. (2022) processed two populations of maize single crosses – one by crossing DHs from two groups and the other a biparental population. The additive-dominance with epistasis model provided generally comparable to slightly higher prediction accuracies across traits and environments, especially for grain yield and the biparental population. Using a multiple-hybrid population of maize, Sang *et al*. (2022) demonstrated that epistasis strongly affected heterosis, especially in the temperate x tropical population. Predicting heterosis for grain yield, they observed an increase of 6.5% in the accuracy by fitting the additive x additive effects, relative to the additive-dominance model. Regarding the impact of modelling epistasis on the breeding value prediction accuracy in METs of DH/inbred/pure lines, we can state that an increase in the prediction accuracy depends on trait, i.e., the contribution of the epistatic effects for the genotypic value or genotypic variance, heritability, and correlation between environments (Vojgani *et al*. 2021; Raffo *et al*. 2022; Vojgani *et al*. 2023).

We designed this simulation-based study to provide theoretical information on the influence of epistasis on genomic prediction efficiency in maize hybrid breeding. The objective was to assess the impact of epistasis on the genomic prediction of non-assessed and non-overlapping single crosses in METs, assuming seven types of digenic epistasis.

## Materials and methods

For simulating the genome, the DH lines from two heterotic groups, and the single crosses, we used the software *REALbreeding* (available by request). The software simulates individual genotypes for genes and molecular markers and phenotypes in three stages, using inputs from the user. The first stage is genome simulation. In this step, the user defines the number of chromosomes, type, number, and density of markers, number of traits, and number of genes (QTLs – major genes and minor genes) for each trait. The second stage is population simulation. In this step, the user specifies the average frequencies for genes and markers, type – full-sib, half-sib, or inbred – and size of progeny, number of random cross or selfing generations, and number of simulations (resampling of the population). The last stage is trait simulation. In this step, the user specifies the maximum and minimum genotypic values for homozygotes (assuming no epistasis), maximum and minimum phenotypic values – to avoid outliers, degree and direction of dominance, and broad sense heritability. The software was used in several studies published since 2013, involving genomic selection/genomic prediction of complex traits, genome-wide association study, and QTL mapping. The software has GUI and versions for Windows, Linux, and MacOS. It can be used for studies in human genetics, plant and animal genetics and breeding, population genetics, and evolution. The current version allows digenic epistasis, pleiotropy, and genotype x environment interaction. It allows recurrent selection and diallel crosses for panmictic populations and DH/inbred/pure lines. Its most significant feature is a strong quantitative genetics background. The additive, dominance, and epistatic genetic values, general and specific combining ability effects, or genotypic values, depending on the population, are computed and not sampled from a probability distribution.

The markers and gene frequencies are generated using beta distribution. The reference population (generation 0) in linkage disequilibrium (LD) is generated by crossing two populations in linkage equilibrium, followed by a generation of random crosses to achieve Hardy-Weinberg equilibrium. The LD value in the gametic pool of generation −1 is 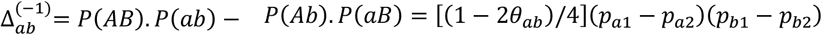, where *θ* is the recombination rate, *p* is the allele frequency, and the indexes 1 and 2 refer to the parental populations. From the maximum and minimum genotypic values for homozygotes, inputted from the user, *REALbreeding* computes the parameter **a** – deviation between the genotypic value for the homozygote of greater expression and the mean of the homozygotes (**m**) – for each gene. The inputs for the degree and direction of dominance allow computing the dominance deviation (**d**) for each gene. Regarding the computation of the nine epistatic effects (I22, I21, I20, *…*, and I00) for each pair of interacting genes, the software initially samples a random value for I22 (the epistatic value for the genotype AABB). This is necessary because there are nine genotypes, nine epistatic effects but only two, three, or four genotypic values, depending on the digenic epistasis. Then, it solves the system of equations for computing the other eight epistatic effects, respecting that each genotypic value should be positive and meet the epistasis type. For example, assuming complementary epistasis, the genotypic values should meet G22 = G21 = G12 = G11 and G20 = G10 = G02 = G01 = G00. To get the I22 random values for each pair of interacting genes, the user should specify the ratio epistatic variance/sum of additive and dominance variances. Computation of the additive x additive (AxA), additive x dominance (AxD), dominance x additive (DxA), and dominance x dominance (DxD) epistatic genetic values is based on Kempthorne (1954). Thus, the software uses the allele frequencies, the LD values, the parameters **m, a**, and **d** for each gene, and the epistatic values for each pair of interacting genes to compute genetic and genotypic values. Finally, using the broad sense heritability, it samples random errors and computes the phenotypic values.

In the first step, we defined 10 chromosomes of 200 cM, 60 K SNPs, and 500 genes, equally distributed on the chromosomes, at random positions. The average density for SNPs and genes were 0.03 and 4 cM, respectively. In the second step, we instructed *REALbreeding* to sample 70 DHs from each heterotic group and to cross the DHs in a partial diallel for generating 4900 single crosses. The average maf for SNPs were 0.25 and 0.20 in the heterotic groups. The average frequency for the favorable allele was 0.35 in one group and 0.50 in the other group. Additionally, we defined 50% of epistatic genes, i.e., 125 pairs of epistatic genes, assuming complementary (9:7 in F_2_), duplicate (15:1 in F_2_), dominant (12:3:1 in F_2_), recessive (9:3:4 in F_2_), and dominant and recessive (13:3 in F_2_) epistasis, duplicate genes with cumulative effects (9:6:1 in F_2_), and non-epistatic gene interaction (9:3:3:4 in F_2_). We also assumed all types of epistasis. In this case, *REALbreeding* took a type at random for each pair of interacting genes. Finally, we instructed the software to allocate all single crosses in five environments. The assumptions regarding environment and genotype x environment effects are described by David *et al*. (2023). We emphasize that these effects are attributable for each gene. In the last step, we defined positive dominance with degree of dominance in the range 0.0 to 1.2 (0.6 on average). The minimum and maximum genotypic values were 50 and 240 g/plant and the minimum and maximum phenotypic values were 30 and 300 g/plant. Finally, we defined heritability at the entry level in the range 30 to 90% over the environments. Note that we did not request *REALbreeding* to simulate, let’s say, r replications in each environment with n plants/plot. Assuming this, the single cross phenotype file would have r x n x 5 × 4900 rows (for example, assuming r = 4 and n = 20, 1960000 rows).

Regarding the distribution of the single crosses over the environments, we adopted 10 and 30% of tested single crosses (490 and 1470, respectively) and the field sparse testing system recommended by Jarquin *et al*. (2020): a large proportion of non-overlapping single crosses (80%) and a small proportion of overlapping single crosses (20%) in the environments. The sampling of single crosses for testing was replicated 30 times. The linear mixed model was *y* = *Xβ* + *Z*_1_*g*_1_ + *Z*_2_*g*_2_ + *Z*_3_*s* + *Z*_4_*aa* + *Z*_5_*ad* + *Z*_6_*dd* + *Z*_7_*g*_1_*e* + *Z*_8_*g*_2_*e* + *Z*_9_*se* + *Z*_10_*aae* + *Z*_11_*ade* + *Z*_12_*dde* + ε. The assumptions for the vectors of general combining ability effects and the vector of specific combining ability effects were 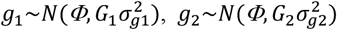 and 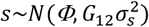, where *G*_1_and *G*_2_ are the additive genomic relationship matrices for the DH lines and *G*_12_ is the dominance genomic relationship matrix (Massman *et al*. 2013). We adopted VanRaden (2008)’s first method for the additive genomic relationship matrix.

Su *et al*. (2012), Vitezica *et al*. (2013), and Nishio and Satoh (2014) derived dominance genomic relationship matrix assuming individuals in a population in Hardy-Weinberg equilibrium. Because its computation from an R package requires the SNP genotypes for the individuals, a simple way of using any dominance genomic relationship matrix is to generate the SNP genotypes for the single crosses, using the DH lines genotypes. Based on Cockerham (1961), Melchinger *et al*. (2023) defined *G*_12_ = *G*_1_*⨂G*_2_. We used an alternative approach, based on the probability of two single crosses having two pairs of alleles identical by descent or identical by state, due to LD between SNPs and genes. That is, we used products of elements from the additive genomic relationship matrix for the DHs of the two heterotic groups. Define *G*_*A*_ = {*p*_*ij*_} (*i, j* = 1 *to n*1 + *n*2) as the additive genomic relationship matrix for the DH lines from the two heterotic groups, where *p*_*ij*_ is the probability of two DHs having alleles identical by descent or identical by state, due to LD between SNPs and genes. Assuming no LD between SNPs and genes, *p*_*ij*_ = *r*_*ij*_, where *r*_*ij*_ is the coefficient of relationship between the DHs. Thus, the dominance genomic relationship matrix for the single crosses is *G*_*D*_ = {*p*_(*ij*)(*kl*)_} = {*p*_*ik*_. *p*_*jl*_ + *p*_*il*_. *p*_*jk*_} (*i, j, k, l* = 1 *to n*1 + *n*2), where *p*_(*ij*)(*kl*)_ is the probability of two single crosses having two pairs of alleles identical by descent or identical by state, due to LD between SNPs and genes. Assuming no LD between SNPs and genes, *p*_(*ij*)(*kl*)_ = *r*_*ik*_. *r*_*jl*_ + *r*_*il*_. *r*_*jk*_.

As for the additive and dominance genomic relationship matrices, the derivation of the epistatic genomic relationship matrices also assumes individuals in a population in Hardy-Weinberg equilibrium (Vitezica *et al*. 2017). We assumed the following assumptions for the vectors of single crosses epistatic genetic values: 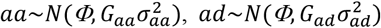, and 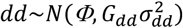, where *ad* = *ad* + *da* and 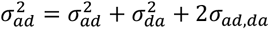. Following Vitezica *et al*. (2017), we defined the AxA, AxD, and DxD genomic relationship matrices as *G*_*aa*_ = *G*_*a*_•*G*_*a*_/[*tr*(*G*_*a*_•*G*_*a*_)/*n*], *G*_*ad*_ = *G*_*a*_•*G*_*D*_/[*tr*(*G*_*a*_•*G*_*D*_)/*n*], and *G*_*dd*_ = *G*_*D*_•*G*_*D*_/[*tr*(*G*_*D*_•*G*_*D*_)/*n*], where • indicates the Hadamard product and *G*_*a*_ is the additive genomic relationship matrix for the single crosses. It can be easily calculated by an R package using the SNP genotypes for the single crosses. We used elements of *G*_*A*_ since *G*_*a*_ = {*q*_(*ij*)(*kl*)_} = {(1/4)(*p*_*ik*_ + *p*_*il*_ + *p*_*jk*_ + *p*_*jl*_)}, where *q*_(*ij*)(*kl*)_ is the probability that two single crosses having alleles identical by descent or identical by state, due to LD between SNPs and genes. Assuming no LD between SNPs and genes, *q*_(*ij*)(*kl*)_ = (1/4)(*r*_*ik*_ + *r*_*il*_ + *r*_*jk*_ + *r*_*jl*_).

Regarding the vectors of interaction with environments and of error effects, to avoid over parametrization, we assumed 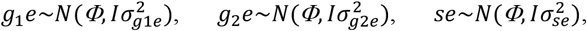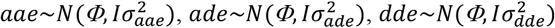, and *ε*∼*N*(*Φ, Iσ*^2^). The simulated dataset was processed using BGLR (de los Campos and Rodriguez 2024). We used the defaults for burn in (500) and iterations (2000). To assess the efficacy of the genomic prediction of tested, untested, and non-overlapping single crosses under epistasis, ignoring and fitting epistasis, we computed the coincidence index and the prediction accuracy, in each environment and across environments. The coincidence index expresses the fraction of n superior predicted single crosses that are among the n superior single crosses. We assumed n = 490. As a reference, we computed the maximum coincidence index, assuming the phenotyping of all single crosses. The true genotypic values for the single crosses are provided by *REALbreeding* (*Gij* = μ + *gi* + *gj* + *sij* + *Iij*). The software also allows getting the true genotypic value in each environment since it provides the true interaction effects g*ije*(*k*). The predicted genotypic value in the environment k is 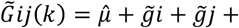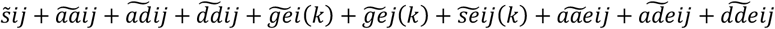 and the predicted genotypic value across environments is 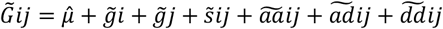. The prediction accuracy was computed as the Pearson’s correlation between the true and the predicted genotypic values, considering only the untested and non-overlapping single crosses. For the scenarios of lower and higher contribution of the epistatic effects to the single cross genotypic variance, we also processed the dataset ignoring epistasis (additive-dominance model).

### Results

The numbers of fixed genes in the two groups of DHs were 15 and 0 (data not shown). Using the complete dataset, the correlations between the heritability and the environmental index ranged from −0.04, with duplicate genes with cumulative effects, to 0.09, assuming dominant and recessive epistasis (0.04 on average). Irrespective of the epistasis type, the heritabilities over the environments ranged from 0.06 to 0.89 (Table 1). The values were higher for dominant and recessive and recessive epistasis and lower for duplicate genes with cumulative effects and non-epistatic gene interaction. There were contrasting environments since the range for the environmental index was approximately 14 g/plant (1120 kg/ha assuming 80000 plants), also regardless of the epistasis type. The percentage of the genotypic variance due to epistasis ranged from 18 to 48%, assuming complementary and duplicate epistasis, respectively. Using the genotypic values for the DHs and single crosses (data not shown), the heterosis ranged from 10 to 25%, under dominant and recessive epistasis and duplicate genes with cumulative effects, respectively.

**Table 1.**
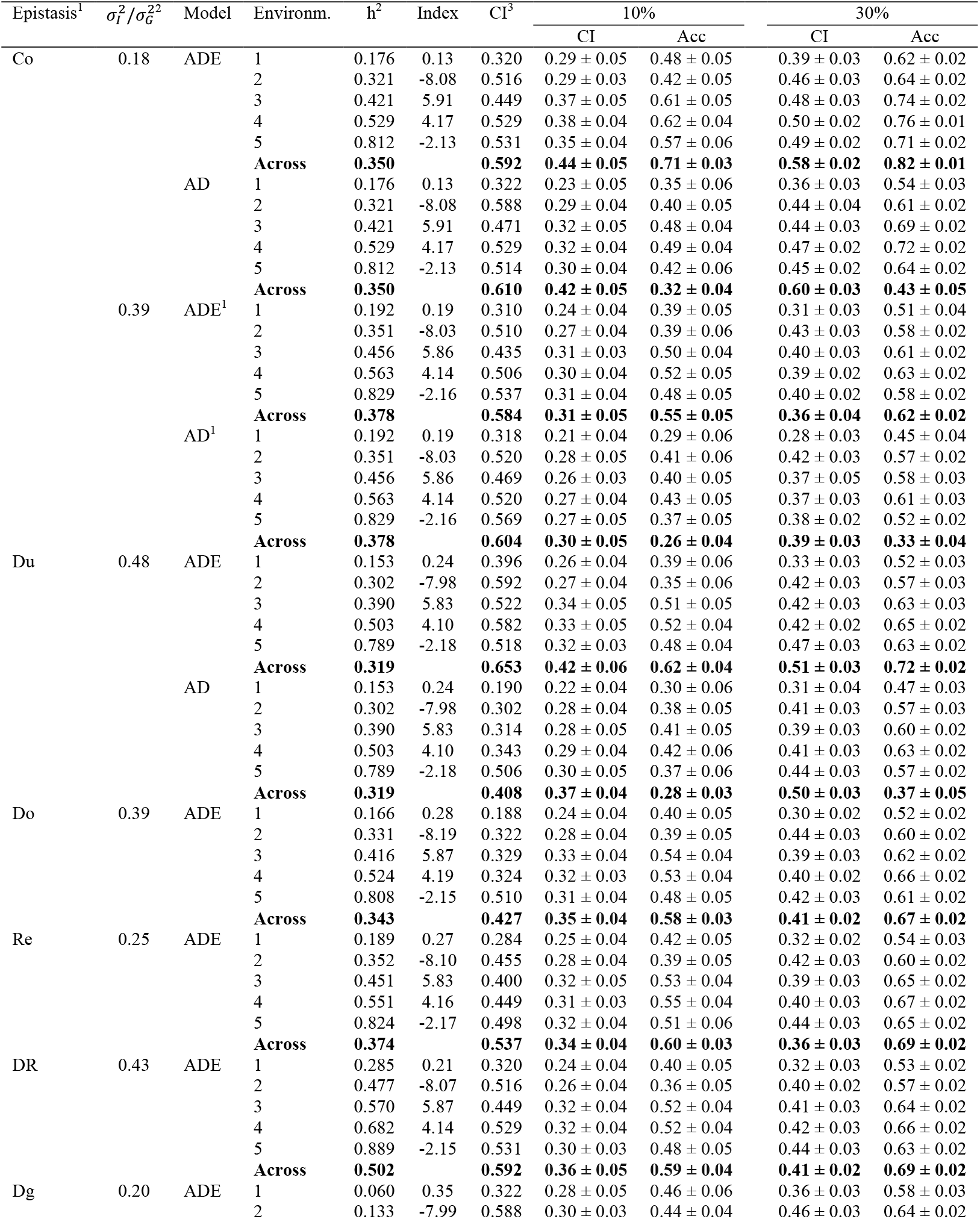
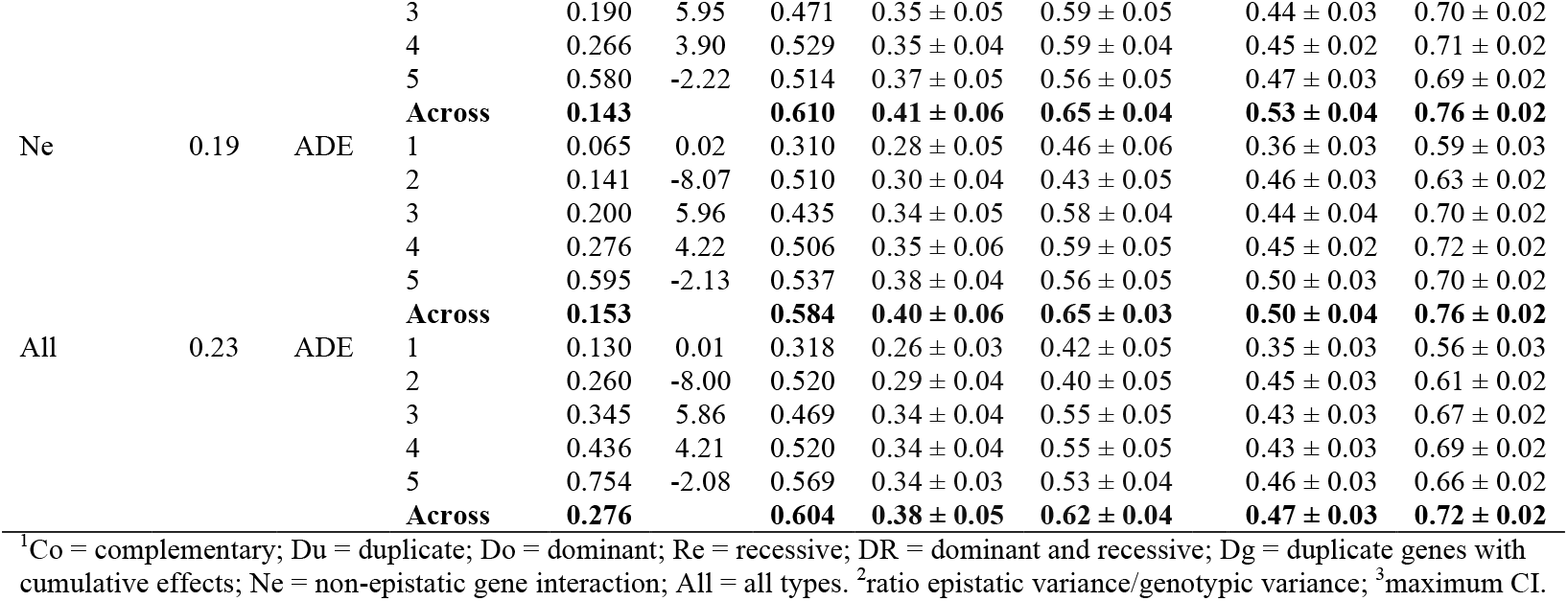
Coincidence indexes (CI) and prediction accuracies (Acc) for the untested single crosses in each environment and across environments, assuming 10 and 30% of tested single crosses, seven digenic epistasis and an admixture of types, fitting the complete (ADE) and the additive-dominance (AD) models.

Considering that the correlations between the heritability and environmental index for distinct epistasis types are approximately 1.0 (0.98-1.00), the differences between the coincidence indexes and the accuracies are due to the magnitude of the genotypic variance attributable to the epistatic effects. The coincidence index ranged from 0.24 to 0.38, under the lower training set, and between 0.30 to 0.50, assuming the higher training set (Table 1). Increasing the training set increased the coincidence index by 6 to 59% (33% on average). Fitting regression models by environment (or heritability, since the heritability increases from environment 1 to environment 5) for the coincidence index in function of the ratio epistatic variance/genotypic variance, we generally observed a decrease in the coincidence index by increasing the ratio up to 0.3, but an increase in the coincidence index by increasing the ratio above 0.4, regardless of the heritability and the training set size.

Fixing complementary epistasis and increasing the ratio epistatic variance/genotypic variance from 18 to 39% led to a decrease in the coincidence index in the range 6.5 to 22.0%, regardless of the training set (Table 1). The decreases were not correlated with the heritability by environment (correlations of approximately 0.1 and −0.2 for training sets 10 and 30%) but negatively correlated with the environmental index (correlations of approximately −0.9 and −0.8 for training sets 10 and 30%). It is important to emphasize that the coincidence index showed positive correlation with the heritability by environment, ranging from 0.49, assuming complementary epistasis, ratio epistatic variance/genotypic variance of 39%, and 30% of tested single crosses, to 0.91, assuming non-epistatic gene interaction and training set of 10%, and dominant and recessive epistasis and training set of 30% (Table 1 and Figure 2). Compared to the maximum values, the relative coincidence indexes ranged from 0.46 to 1.28 (0.70 on average), under the lower training set, and between 0.67 and 1.63 (0.96 on average), assuming the higher training set, depending on the heritability, epistasis type and ratio epistatic variance/genotypic variance. In general, the coincidence index assuming higher training set can be superior to the maximum coefficient only under low heritability and low proportion of the genotypic variance explained by the epistatic effects.

**Figure 1.**
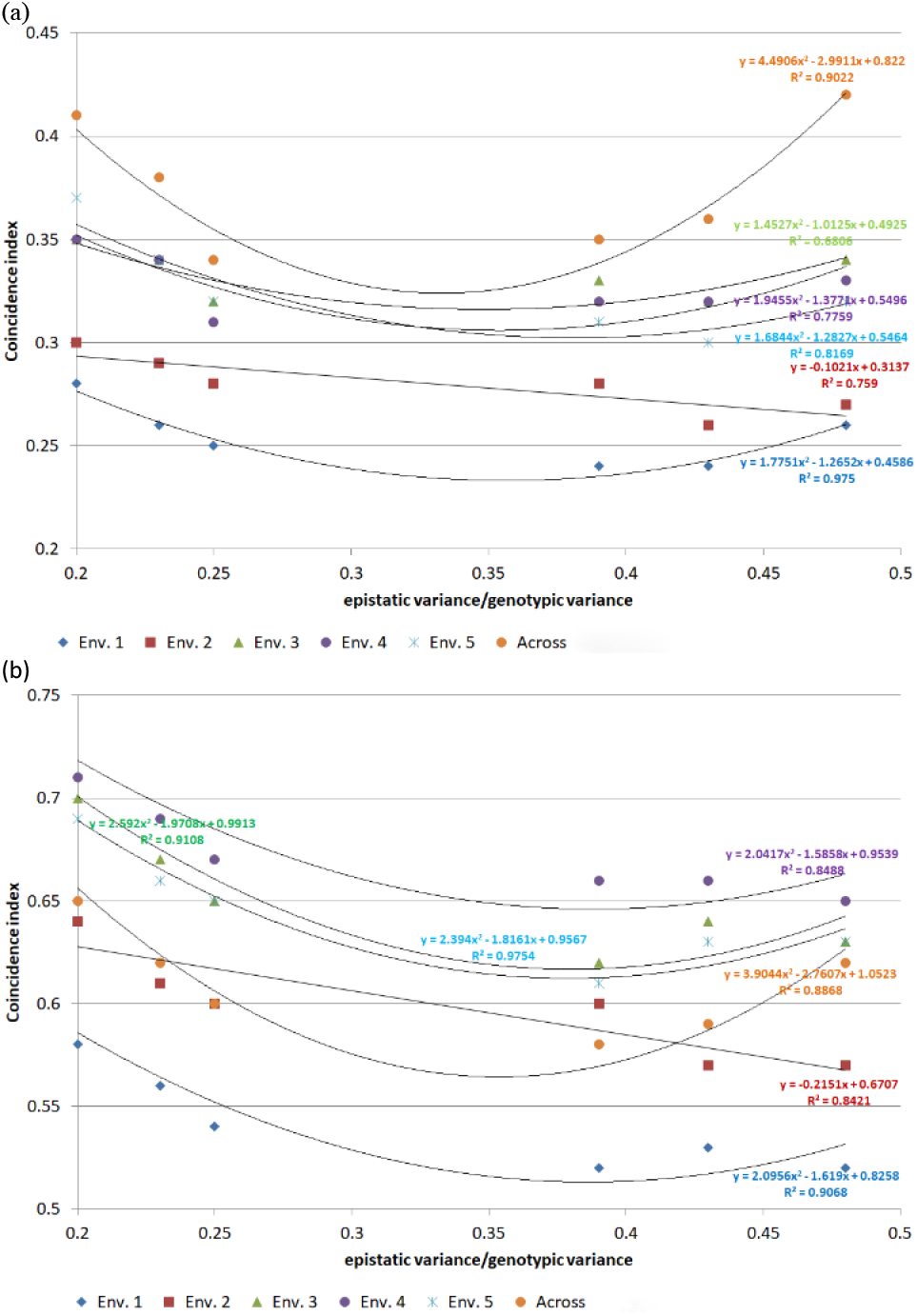
Relationship between the coincidence index and the ratio epistatic variance/genotypic variance by environment (ordered by the average heritability, from low (Env. 1; h^2^ = 15%) to high (Env. 5; h^2^ = 76%)) and across environments (h^2^ = 31%), assuming 10 (a) and 30% (b) of tested single crosses.

**Figure 2.**
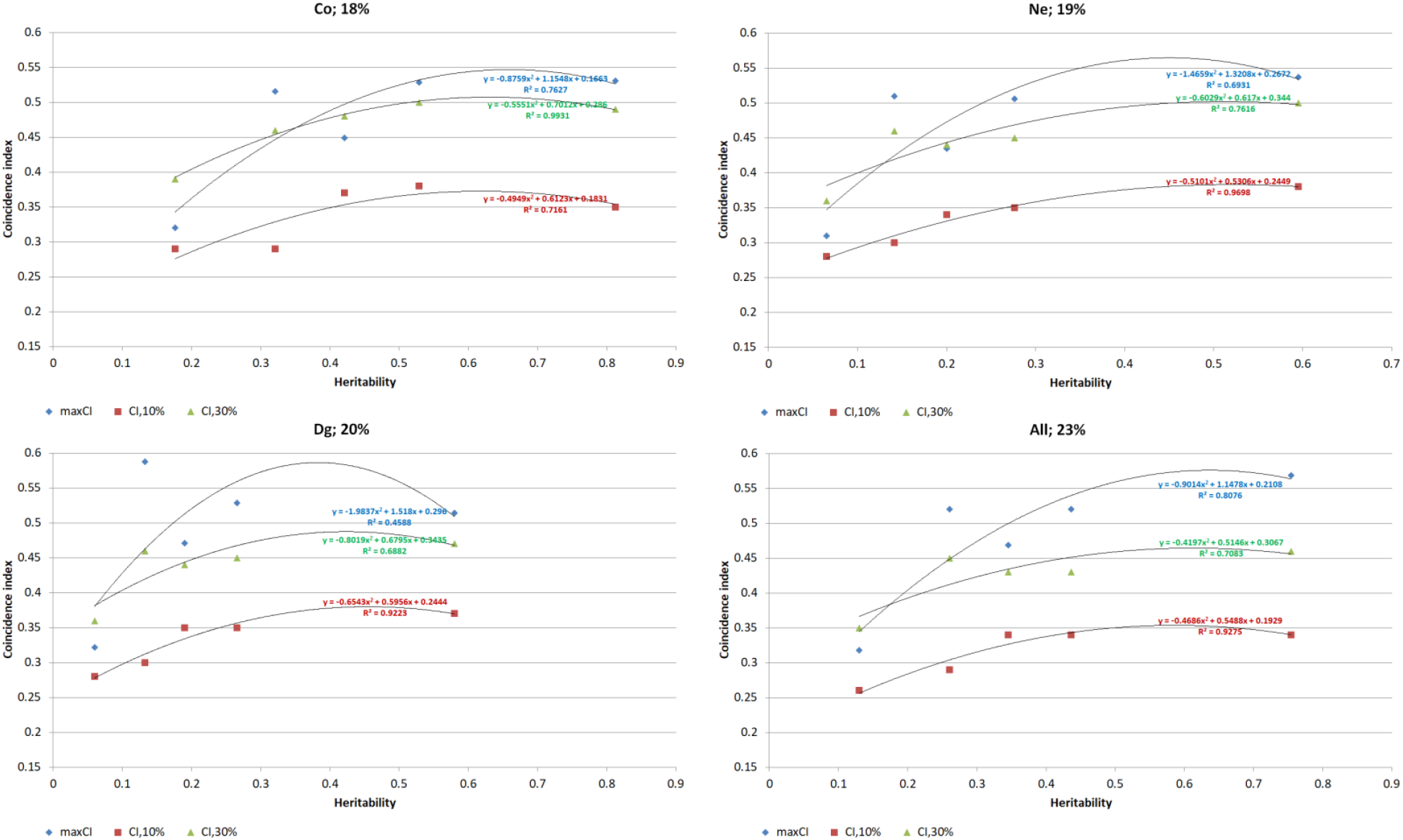
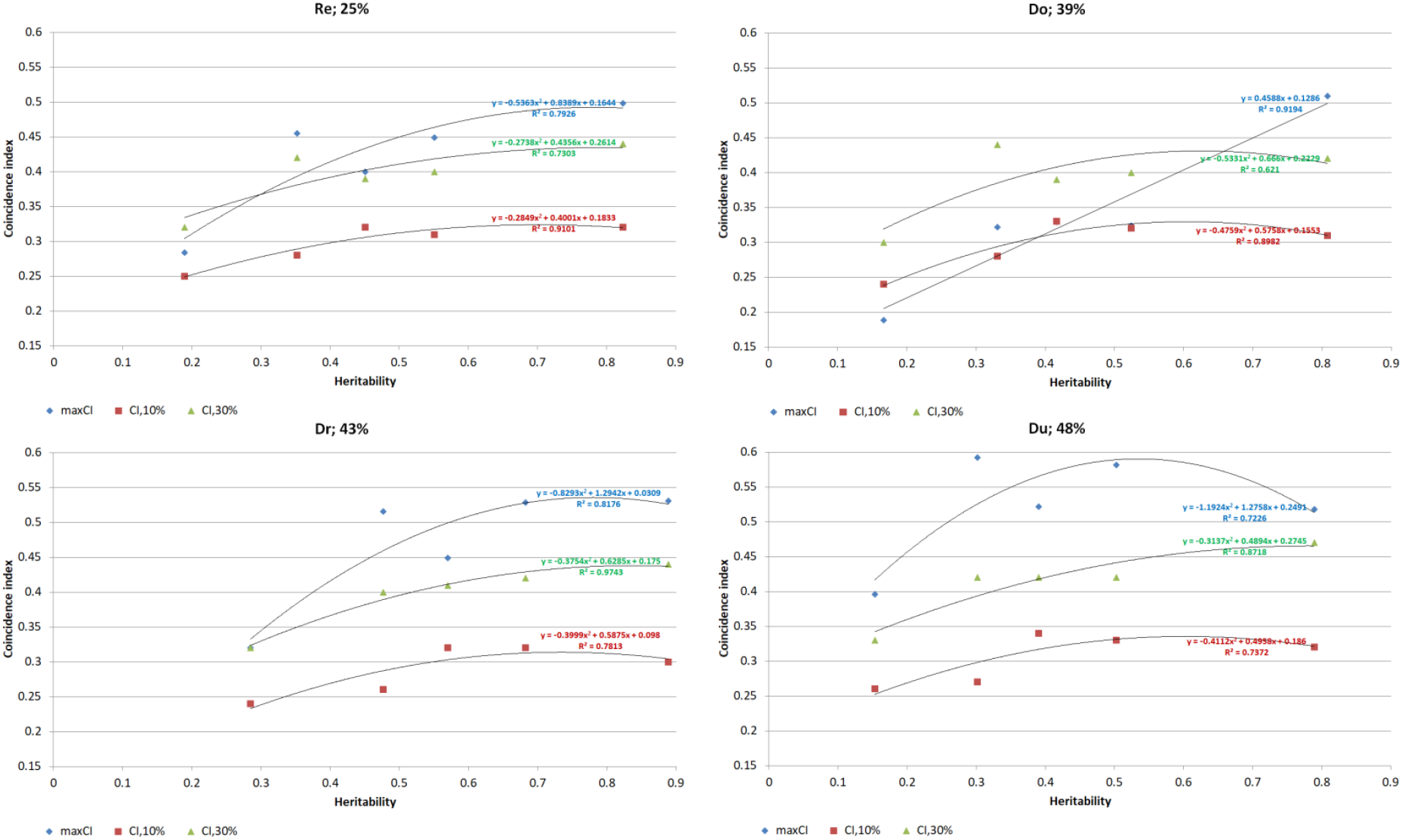
Relationship between the coincidence index and the heritability by environment, assuming phenotyping of all single crosses (maxCI) and training sets of 10 and 30%, for distinct epistasis types and ratios epistatic variance/genotypic variance. Co = complementary; Du = duplicate; Do = dominant; Re = recessive; DR = dominant and recessive; Dg = duplicate genes with cumulative effects; Ne = non-epistatic gene interaction; All = all types.

Furthermore, accuracy showed positive correlation with coincidence index, especially for the lower training set. Under the lower training set, the correlations ranged from 0.82, assuming recessive epistasis and non-epistatic gene interaction, to 0.97, assuming complementary epistasis and ratio epistatic variance/genotypic variance of 18% (Table 1). Increasing the training set decreased the correlations from 8 to 25%, assuming dominant and recessive epistasis and an admixture of types, respectively (−15% on average). The correlations assuming the higher training set ranged from 0.69, assuming all types, to 0.85, assuming dominant and recessive epistasis.

Assuming epistasis but fitting the additive-dominance model led to a decrease in the coincidence indexes, especially under the lower training set (Table 1). Under the lower training set, the coincidence indexes changed from −21 to 4% (−11% on average). Under the higher training set, the coincidence indexes decreased from 2 to 10% (−6% on average). The changes were positively correlated with the heritability by environment (0.47 to 0.83) and, in general, negatively correlated with the environmental index (0.10 to −0.92), especially under the lower training set.

## Discussion

Breeders agree that epistasis can contribute for the genotypic value of a quantitative trait, but the most important component of the genotypic value is the additive. Most of the evidence of epistasis came from studies of inheritance of qualitative traits, QTL mapping, and, in recent years, from GWAS and transcriptome, proteome, and metabolome analyses (Mackay 2014; Domingo *et al*. 2019). In our study, the epistatic effects explained 18 to 48% of the single cross genotypic variance. Some studies using genomic prediction provided evidence that fitting epistasis can significantly improve prediction accuracy (Calleja-Rodriguez *et al*. 2021; Vojgani *et al*. 2021; Vojgani *et al*. 2023). In other investigations the results were inconclusive or evidenced no increase in the prediction accuracy (Vitezica *et al*. 2018; Onogi *et al*. 2021; Alves *et al*. 2023). Because a lower contribution of epistasis to the phenotypic variance (0 to 10%), Bhuiyan *et al*. (2024) observed increases of only 2 to 3% in the prediction accuracy when epistatic effects were included in the model. Using wheat and maize datasets, Jiang and Reif (2015) concluded that prediction accuracy can be improved by modeling epistasis for selfing species but may not for outcrossing species. Our results showed increases from 0 to 37%, depending on the heritability, epistasis type, and ratio epistatic variance/genotypic variance, especially under the lower training set.

Regarding the prediction accuracy of untested maize single crosses in METs, this study agrees with the only two field-based studies modelling epistatic effects. We showed that prediction accuracy is affected by heritability, epistasis type, ratio epistatic variance/genotypic variance, and training set. In the study of GonzÁlez-DiÉguez *et al*. (2021), for grain yield, the epistasis explained only 4% of the broad sense heritability. The prediction accuracies for the untested single crosses assuming the complete model were equal to the values observed for the additive-dominance model. Roth *et al*. (2022) processed two groups of single crosses in METs. Only in the biparental population the AxA effects contributed to grain yield. The AxA effect contributed 33% for the broad sense heritability. Even with this significant contribution, the difference between the average prediction accuracies for the complete and the additive-dominance model was only 0.02 (0.63 and 0.65). Our prediction accuracies of non-assessed and non-overlapping single crosses ranged from 0.35 to 0.76, proportional to the training set size.

Based on the coincidence indexes, we can state that genomic prediction of single crosses in METs is a very effective process, even assuming only 10% of tested single crosses. Assessing 10% of the single crosses, assuming 20% of overlapping single crosses, means a reduction of 96% in the cost to phenotype all single crosses (no genotyping). For sure, then, the overall cost of genotyping the DHs and assessing a sample of the single crosses is lower than the cost to phenotype all single crosses. Depending on the heritability, the epistasis type, and the ratio epistatic variance/genotypic variance, we identified from 24 to 38% of the genetically superior single crosses in each environment. Regarding the identification of the genetically superior single crosses (across environments), the efficacy ranged from 31 to 44%. A very impressive result. Significant increases in efficacy can be achieved by increasing the training set size: 25 to 32% in each environment by increasing the training set size to 30%. The same conclusion was stated for testcrosses (prediction of general combining ability effect) and single crosses in the simulation-based studies of Viana *et al*. (2019) and Viana *et al*. (2018), assuming the additive-dominance model. Their coincidence indexes for single crosses across environments, for the SNP density of 0.1 cM and training set of 10%, ranged from 7 to 65%, depending on the population used for sampling the DHs and heritability. The values were higher for the biparental population – because higher LD – and proportional to the heritability. Increasing the training set to 30% increased the efficacy of identification of the superior single crosses from 15 to 83%. This confirms that epistasis can negatively affects the genomic prediction of single crosses, depending on the epistasis type and the proportion of the genotypic variance due to non-allelic interaction.

Finally, we emphasize that breeders can be confident in the estimated prediction accuracy to assess the efficacy of identification of the superior single crosses in each environment and across environment, since the prediction accuracy is highly correlated with the coincidence index, especially under no epistasis (Viana *et al*. 2018; Viana *et al*. 2019). It is important to emphasize too the role of LD in the genomic prediction of untested single crosses. The high coincidence indexes that we observed are due only to the genomic similarity among the DHs in each group, i.e., LD between SNPs and genes. There was no genetic relationship between the DHs in each group. Based on the coincidence indexes obtained by Viana *et al*. (2018), including genetic relationship between the DHs from S_0_ plants in each group increased from 13 to 26% the efficacy of identification of untested single crosses. However, the increment was inversely proportional to the heritability.

## Conclusions

Assuming a substantial contribution of the epistatic variance to the single cross genotypic variance and fitting the GxE effects, we can conclude: as expected, the heritability in each environment and the training set size significantly affects the efficacy of genomic prediction of untested single crosses; the proportion of the genotypic variance explained by the epistatic variance and the epistasis type also significantly affects the efficacy of genomic prediction of untested single crosses; ignoring the epistatic effects in the analysis can negatively affect the efficacy of genomic prediction of untested single crosses, especially under a low training set; the prediction accuracy efficiently express the efficacy of genomic prediction of untested single crosses; the genomic prediction of untested single crosses in each environment and across environments is very effective, even under a low training set size.

## Acknowledgments

We thank the National Council for Scientific and Technological Development (CNPq), the Brazilian Federal Agency for Support and Evaluation of Graduate Education (CAPES; Finance Code 001), and the Foundation for Research Support of Minas Gerais State (FAPEMIG) for financial support.

## Author contributions

**JMSV** Conceptualization, Software, Funding acquisition, Data curation, and Writing - original draft. **JPAS** Formal analysis and Writing – review.

## Data availability

The dataset is available at https://doi.org/10.6084/m9.figshare.27063610.v2.

